# Comparative analysis of antigen-specific anti-SARS-CoV-2 antibody isotypes in COVID-19 patients

**DOI:** 10.1101/2020.12.04.407510

**Authors:** Hidetsugu Fujigaki, Masato Inaba, Michiko Osawa, Saya Moriyama, Yoshimasa Takahashi, Tadaki Suzuki, Kenya Yamase, Yukihiro Yoshida, Yo Yagura, Takayoshi Oyamada, Masao Takemura, Yohei Doi, Kuniaki Saito

## Abstract

Serological tests for detection of anti-severe acute respiratory syndrome coronavirus 2 (SARS-CoV-2) antibodies in blood are expected to identify individuals who have acquired immunity against SARS-CoV-2 and indication of seroprevalence of SARS-CoV-2 infection. Many serological tests have been developed to detect antibodies against SARS-CoV-2. However, these tests have considerable variations in their specificity and sensitivity, and whether they can predict levels of neutralizing activity is yet to be determined. This study aimed to investigate the kinetics and neutralizing activity of various antigen-specific antibody isotypes against SARS-CoV-2 in serum of coronavirus disease 2019 (COVID-19) patients confirmed via polymerase chain reaction test. We developed IgG, IgM and IgA measurement assays for each antigen, including receptor-binding domain (RBD) of spike (S) protein, S1 domain, full length S protein, S trimer and nucleocapsid (N) domain, based on enzyme-linked immunosorbent assay. The assays of the S protein for all isotypes showed high specificity, while the assays for all isotypes against N protein showed lower specificity. The sensitivity of all antigen-specific antibody isotypes depended on the timing of the serum collection and all of them, except for IgM against N protein, reached more than 90% at 15-21 days post-symptom onset. The best correlation with virus neutralizing activity was found for IgG against RBD (RBD-IgG), and levels of RBD-IgG in sera from four severe COVID-19 patients increased concordantly with neutralizing activity. Our results provide valuable information regarding the selection of serological test for seroprevalence and vaccine evaluation studies.

## Introduction

Coronavirus disease 2019 (COVID-19), caused by severe acute respiratory syndrome coronavirus 2 (SARS-CoV-2) infection, has instigated a widespread concern globally (1). As of 1^st^ November 2020, more than 46 million confirmed cases and 1 million deaths have been recorded worldwide. Currently, the gold standard for the diagnosis of COVID-19 is real time RT-PCR assay, which detects SARS-CoV-2 RNA in respiratory tract specimens. In the meantime, serological tests to detect anti-SARS-CoV-2 antibodies in the blood have also been investigated (2, 3). Although the immune response and antibody kinetics against SARS-CoV-2 is not well understood, serological tests are expected to be useful for various purposes (4, 5). For instance, convalescent serum/plasma therapy has been initiated for severe COVID-19 patients in some countries (6, 7). Serological tests that quantify neutralizing antibodies should effectively identify donors with high antibody titers. Serological tests for neutralizing antibodies may also be used to determine the efficacy of vaccines currently being developed (8-10). In addition, serological tests provide insight into an individual’s immune status as well important epidemiological information on spread of the infection in a specific region (2).

To measure an antibody against SARS-CoV-2, it is important to measure an antigen-specific antibody suitable for its application. A typical coronavirus (CoV) contains four major structural proteins: spike (S), membrane (M), envelope (E) and nucleocapsid (N) (11, 12). Previous studies have reported that highly immunogenic antigens of SARS-CoV and Middle East respiratory syndrome coronavirus (MERS-CoV) are S and N proteins (13-16). Therefore, most of the serological assays developed to detect anti-SARS-CoV-2 antibody also target these proteins. The S protein consists of two functional subunits (S1 and S2) which form a homotrimer. S1 contains a receptor-binding domain (RBD) and is required for binding to host cells. Therefore, detection of S1 (RBD)-specific antibody is considered as an index for predicting neutralizing activity that protects host cells from virus infection (17, 18). On the other hand, N protein functions to package the capsid-formed genome into virions and acts as a viral RNA silencing suppressor required for viral replication (12). The N protein of CoV is found to be overexpressed in large quantities during infection and is highly immunogenic. Therefore, antibodies against the N protein are promising indicators for detecting SARS-CoV-2 infection, especially once SARS-CoV-2 vaccines are widely deployed, all of which currently uses whole or partial S protein as the antigen.

Till date, based on S and N proteins, many serological diagnostic methods have been developed to detect antibodies against SARS-CoV-2. However, detailed analysis of antibody isotypes against each antigen, including the S trimer, has not been conducted. Therefore, this study aimed to investigate the kinetics and neutralizing activity of various antigen-specific antibodies against SARS-CoV-2 in the serum of polymerase chain reaction (PCR)-confirmed COVID-19 patients. To assess the various antigen-specific antibodies, we developed the IgG, IgM and IgA measurement assays for each antigen, including RBD, S1, S full length (S full), S trimer and N based on enzyme-linked immunosorbent assay (ELISA). Using this method, we systematically analysed the antibodies in the serum of COVID-19 patients. We also examined the correlation between each antibody and the neutralizing activity in order to clarify which antibody best predicted neutralizing activity. This study advances the understanding of immune response and antibody kinetics against SARS-CoV-2 and informs several applications such as identifying donors with high antibody titers for convalescent serum/plasma therapy and measuring the immunogenicity of vaccines being developed.

## Materials and Methods

### Samples

This study was reviewed and approved by the Ethics Committee for Clinical Research of the Center for Research Promotion and Support in Fujita Health University (authorisation number HM19-493 and HM17-341).

We utilized a series of residual serum samples from 41 COVID-19 patients who were admitted to Fujita Health University Hospital from February 28 to May 21, 2020. The demographic and clinical characteristics of the patients are presented in Table 1 and Supplementary Table 1. All patients were confirmed as COVID-19 cases by real time PCR assay of nasopharyngeal swab specimens at the time of or prior to admission. The date of onset was determined as the day when the patients started experiencing COVID-19 symptoms. Severity classifications were made according to the Guidelines of the Treatment and Management of Patients with COVID-19 published by the Infectious Diseases Society of America. Severe COVID-19 patients were defined as those with SpO2 ≤ 94% on room air, or those who require supplemental oxygen. The other patients were defined as non-severe COVID-19.

**Table 1.** Clinical characteristics of COVID-19 patients.

|  |  | Total (n = 41) |  | Severe (n = 6) |  | Non-severe (n = 35) |  |
| --- | --- | --- | --- | --- | --- | --- | --- |
|  |  | number (%) |  | number (%) |  | number (%) |  |
| Age. Median (min-max) |  | 41 | (16–93) | 54 | (31–66) | 53 | (16–93) |
| Sex | Male | 22 | (53.7%) | 3 | (50.0%) | 19 | (54.3%) |
|  | Female | 19 | (46.3%) | 3 | (50.0%) | 16 | (45.7%) |
| Symptoms | Fever | 33 | (80.5%) | 6 | (100.0%) | 27 | (77.1%) |
|  | Cough | 12 | (29.3%) | 3 | (50.0%) | 9 | (25.7%) |
|  | Dyspnoea | 4 | (9.8%) | 4 | (66.7%) | 0 | (0.0%) |
|  | Diarrhoea | 1 | (2.4%) | 1 | (16.7%) | 0 | (0.0%) |
|  | Dysgeusia | 2 | (4.9%) | 0 | (0.0%) | 2 | (5.7%) |
|  | Dysosmia | 4 | (9.8%) | 0 | (0.0%) | 4 | (11.4%) |
|  | Sore throat | 2 | (4.9%) | 0 | (0.0%) | 2 | (5.7%) |
|  | Asymptomatic | 3 | (7.3%) | 0 | (0.0%) | 3 | (8.6%) |

One hundred serum samples obtained from 100 healthy human volunteers (mean age, 47; males, 58 and females, 42), collected before the COVID-19 pandemic (July 2012), were used as negative controls to evaluate the specificity and cut-off values for each assay. All serum samples (aliquoted and stored at −80°C) were thawed and evaluated at the same time for the analyses.

### ELISA development and assay

SARS-CoV-2 RBD, S1, S full, S trimer and N protein expressed in HEK293 cells were selected as the target antigens. 96-well plates (Thermo Fisher Scientific, USA) were coated per well with 250 ng of individual recombinant proteins, covered and incubated overnight at 4°C. The plates were washed with phosphate buffer saline containing 0.1% Tween20 (PBST), blocked with PBST containing 1% casein protein stored overnight at 4°C. After blocking, plates were vacuum dried.

Serum samples were heated at 56°C for 30 min before use in order to inactivate any potential residual virus in the sera. Serum samples were diluted by 1:201 for IgG and IgA assays or 1:2010 for IgM assay in sample buffer containing 2% bovine serum albumin. Afterwards, 50 μL of the diluted serum samples were added per well and incubated at room temperature (RT) for 60 min. The plates were washed three times with 300 μL PBST and 50 μL of peroxidase-labelled anti-human IgG, IgM or IgA antibody was added per well and incubated at RT for 60 min. After incubation, the plates were washed five times with 300 μL PBST and 100 μL of substrate solution (TMB/H_2_O_2_) was added per well and incubated at RT for 10 min. Subsequently, the reaction was stopped by addition of 100 μL of 1M HCl. The absorbance was measured at 450 nm using a 620 nm reference filter in a microplate reader (Thermo Fisher Scientific, USA). The antibody titers were expressed as U/mL.

### Virus neutralizing assays

Serum samples were heat-inactivated at 56°C for 30 minutes and were serially diluted with Dulbecco modified Eagle medium (Fujifilm Wako Pure Chemical, Japan) supplemented with 2% fetal bovine serum (Biowest, USA) and 100 unit/mL penicillin, 100 μg/mL streptomycin (Thermo Fisher Scientific, USA). The mixture of diluted sera and 100 TCID_50_ SARS-CoV-2 JPN/TY/WK-521 strain were incubated at 37°C for 1 hour, then placed on VeroE6/TMRRSS2 cells (JCRB1819, JCRB Cell Bank) and cultured at 37°C with 5% CO_2_ (19). On day 5, plates were fixed with 20% formalin (Fujifilm Wako Pure Chemical, Japan) and stained with crystal violet solution (Sigma-Aldrich, USA) for the evaluation of cytopathic effect (CPE). The index of the highest sera dilution factor with CPE inhibition was defined as microneutralization test titer (MNT).

### Statistical analysis

Statistical analysis was performed using GraphPad Prism version 8.0.0 for Windows (GraphPad Software, USA). Sensitivity, specificity, and receiver operating characteristic (ROC) were calculated based on the RT-PCR results. Correlations analysis between antibody titre and neutralization test was performed using Spearman’s correlation coefficient. Statistical difference between non-severe and severe COVID-19 patients was determined using two-tailed Mann-Whitney test and p-value less than 0.05 was considered statistically significant.

## Results

### Antigen-specific antibody isotype responses in COVID-19 patients

We evaluated the ELISA designed to detect SARS-CoV-2 antigen-specific IgG, IgM and IgA against RBD, S1, S full, S trimer and N protein. To quantify the antibody responses to each antigen, we tested 169 serums from 41 SARS-CoV-2-infected patients and 100 negative control serums from healthy donor collected before SARS-CoV-2 pandemic. We categorized sera from SARS-CoV-infected cases according the timing of their collection relative to symptom onset. All antigen-specific antibodies and their isotypes started to increase as early as 0–7 days after the onset of symptoms compared to the negative control samples, which showed no marked response to any of the antigens (Fig. 1). The IgG and IgM levels against all antigens continued to increase over time, while IgA levels peaked by 15–21 days and declined thereafter.

**Figure 1.**
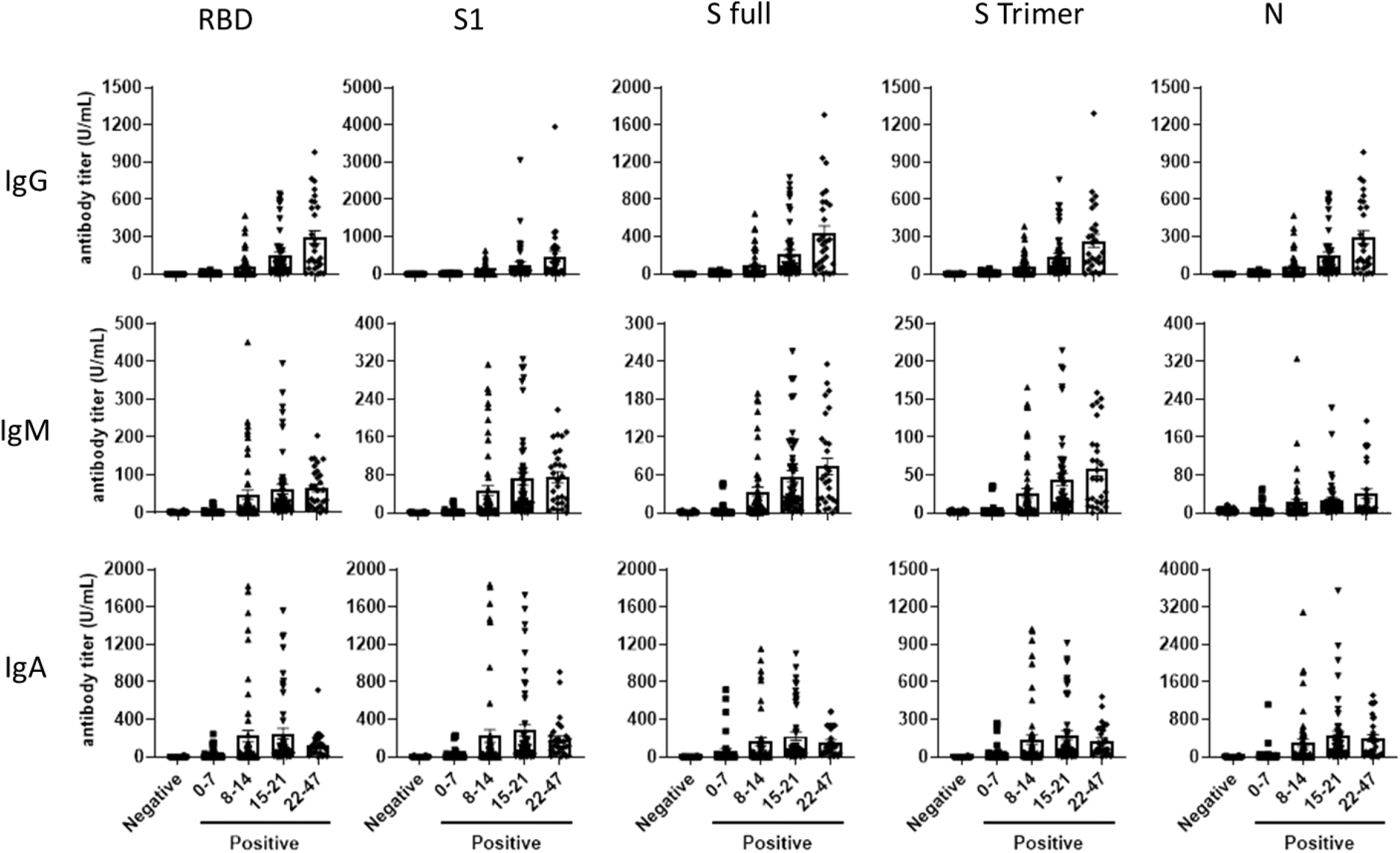
Antibody isotype responses against each SARS-CoV-2 antigen in COVID19 patients. Positive samples from 41 COVID-19 patients (days 0–7 (n = 37), days 8–14 (n = 54), days 15–21 (n = 49) and days 22–47 (n = 29) from symptom onset) and negative samples (n = 100) were tested for IgG, IgM and IgA antibodies against SARS-CoV-2 RBD, S1, S full, S trimer and N antigens using ELISAs. Error bars represent standard errors of the mean.

### Diagnostic performance of the developed ELISAs

To evaluate the diagnostic performance and determine the cut-off values of the developed ELISAs, ROC analysis was performed using the same samples. The ROC analysis showed that the diagnostic performance differed according to the antigens and their isotypes (Fig. 2). The best results were obtained with S1-IgG (area under curve [AUC], 0.943). Conversely, N-IgM showed the lowest diagnostic performance (AUC = 0.797). The analysis also showed that AUC of all antigen-specific antibodies and their isotypes increased over time after the onset of symptoms (Fig. 2).

**Figure 2.**
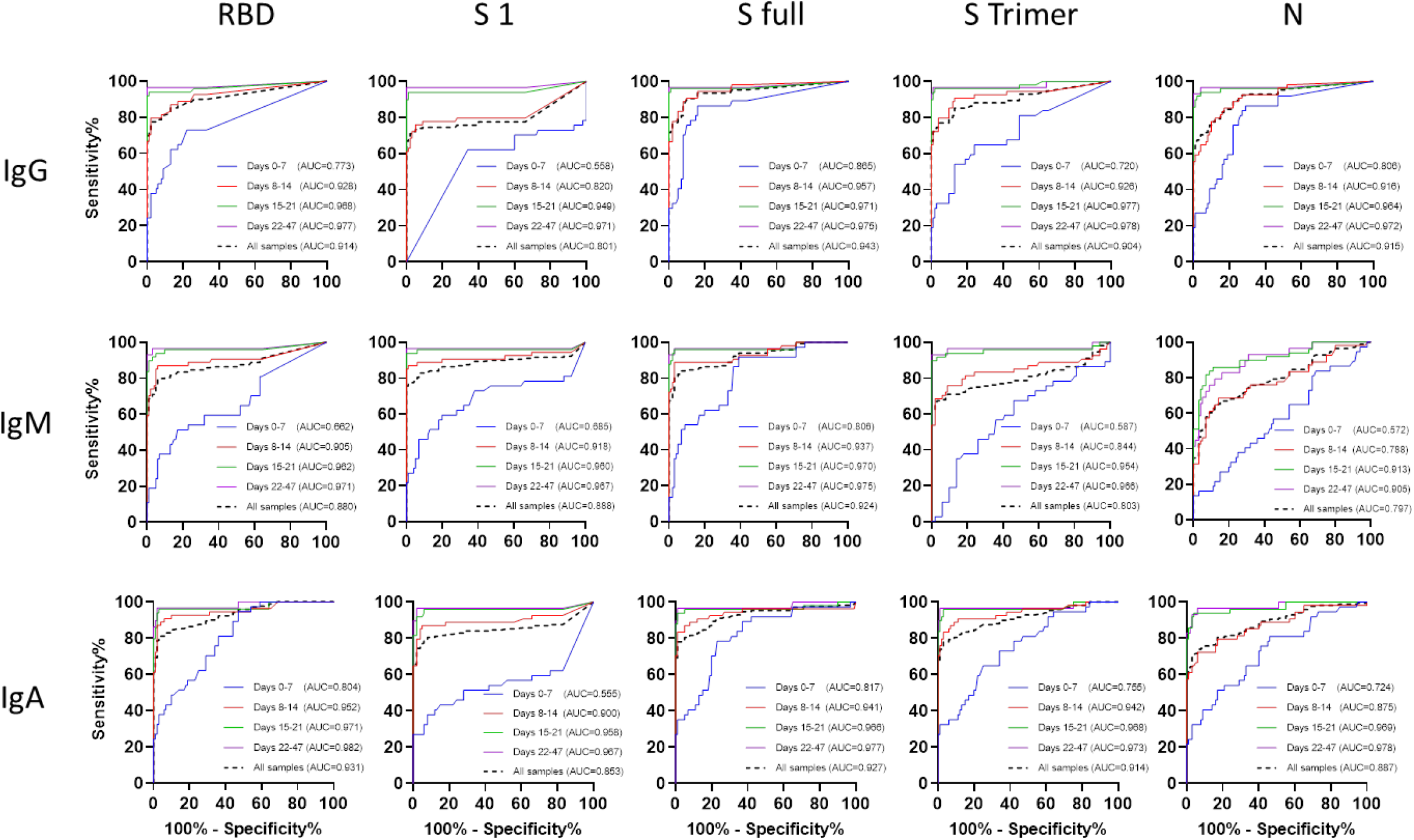
Receiver operating characteristics (ROC) analysis of antibody isotypes for each SARS-CoV-2 antigen over several time periods after symptom onset. COVID-19 negative (n = 100) and positive (n = 169) serum samples from 41 patients (days 0–7 (n = 37), days 8–14 (n = 54), days 15–21 (n = 49) days 22–47 (n = 29) after symptom onset) w ere analysed.

To evaluate the specificity and sensitivity of the developed ELISAs, the optimal cut-off values for each antigen-specific antibody isotype were determined by the Youden’s index using the ROC analysis of all samples (Table 2). The specificity of the S protein (RBD, S1, S full and S Trimer) assays for all isotypes showed comparable results, with high specificity ranging from 94.0%–99.0%. On the other hand, the assays against N protein for all isotypes showed lower specificities of 79.0%, 88.0% and 82.0% for N-IgG, N-IgM and N-IgA, respectively. The sensitivity was dependent on the timing of the serum collection. During 0–7 days after symptom onset, the sensitivity of antigen-specific IgG, IgM and IgA were considerably different and ranged between 23.7%–57.9%, 13.2%–44.7% and 31.6%–50.0%, respectively. Thereafter, the sensitivity of antigen-specific IgG, IgM and IgA increased to more than 90% at 15–21 days after symptom onset, except for N-IgM (85.7%). The sensitivity of all antigen-specific antibody isotypes increased to maximal levels at 22–47 days after symptom onset and ranged between 93.1%–96.6%, except for N-IgM (79.3%).

**Table 2.** Specificity and sensitivity of antibody isotypes for each SARS-CoV-2 antigen based on time periods after symptoms onset.

|  |  |  |  |  |  |  |
| --- | --- | --- | --- | --- | --- | --- |
| **IgG antibodies** |  |  |  |  |  |  |
|  |  | **RBD** | **S1** | **S full** | **S Trimer** | **N** |
|  | **Cut-off value (U/mL)** | 0.86 | 1.10 | 0.70 | 1.80 | 0.70 |
| **Specificity** | **Negative control (n = 100)** | 98.0% | 97.0% | 93.0% | 96.0% | 79.0% |
| **Sensitivity** | 0–7 days (n = 37) | 39.5% | 23.7% | 47.4% | 31.6% | 57.9% |
|  | 8–14 days (n = 54) | 79.6% | 72.2% | 83.3% | 79.6% | 85.2% |
|  | 15–21 days (n = 49) | 93.9% | 93.9% | 95.9% | 95.9% | 95.9% |
|  | 22–47 days (n = 29) | 96.6% | 96.6% | 96.6% | 96.6% | 96.6% |
| **IgM antibodies** |  |  |  |  |  |  |
|  |  | **RBD** | **S1** | **S full** | **S Trimer** | **N** |
|  | **Cut-off value (U/mL)** | 1.70 | 2.07 | 1.10 | 3.12 | 5.65 |
| **Specificity** | **Negative control (n = 100)** | 94.0% | 99.0% | 94.0% | 97.0% | 88.0% |
| **Sensitivity** | 0–7 days (n = 37) | 36.8% | 26.3% | 44.7% | 13.2% | 21.1% |
|  | 8–14 days (n = 54) | 87.0% | 85.2% | 88.9% | 68.5% | 64.8% |
|  | 15–21 days (n = 49) | 93.9% | 93.9% | 95.9% | 91.8% | 85.7% |
|  | 22–47 days (n = 29) | 96.6% | 96.6% | 96.6% | 93.1% | 79.3% |
| **IgA antibodies** |  |  |  |  |  |  |
|  |  | **RBD** | **S1** | **S full** | **S Trimer** | **N** |
|  | **Cut-off value (U/mL)** | 1.85 | 1.89 | 4.37 | 3.40 | 3.00 |
| **Specificity** | **Negative control (n = 100)** | 97.0% | 94.0% | 99.0% | 97.0% | 82.0% |
| **Sensitivity** | 0–7 days (n = 37) | 36.8% | 31.6% | 34.2% | 31.6% | 50.0% |
|  | 8–14 days (n = 54) | 87.0% | 87.0% | 83.3% | 83.3% | 79.6% |
|  | 15–21 days (n = 49) | 95.9% | 95.9% | 93.9% | 95.9% | 93.9% |
|  | 22–47 days (n = 29) | 96.6% | 96.6% | 96.6% | 96.6% | 96.6% |

### Correlation between antigen-specific antibody isotypes and neutralizing activity

Next, we examined the relationship between antigen-specific antibody isotype levels and virus neutralizing activity in sera collected from the COVID-19 patients. We performed neutralization assays in a subset of samples including 34 sera from 19 COVID-19 patients. Fig. 3A shows the correlation between each of the antigen-specific antibody isotypes and the neutralizing activity (MNT). All antigen-specific isotypes, except for N-IgM (Spearman r = 0.150, p-value = 0.397) and N-IgA (Spearman r = 0.172, p-value = 0.332), showed significant positive correlation between antibody levels and neutralizing activity (Fig. 3A). The best correlation was found with RBD-IgG (Spearman r = 0.751, p-value < 0.0001). Fig. 3B presents the Spearman correlation coefficient (r) of each antigen-specific antibody isotypes. Overall, IgG showed good correlation with neutralizing activity in all the antigens. Conversely, IgA showed a relatively lower correlation with neutralizing activity and N-IgA in particular showed no significant correlation.

**Figure 3.**
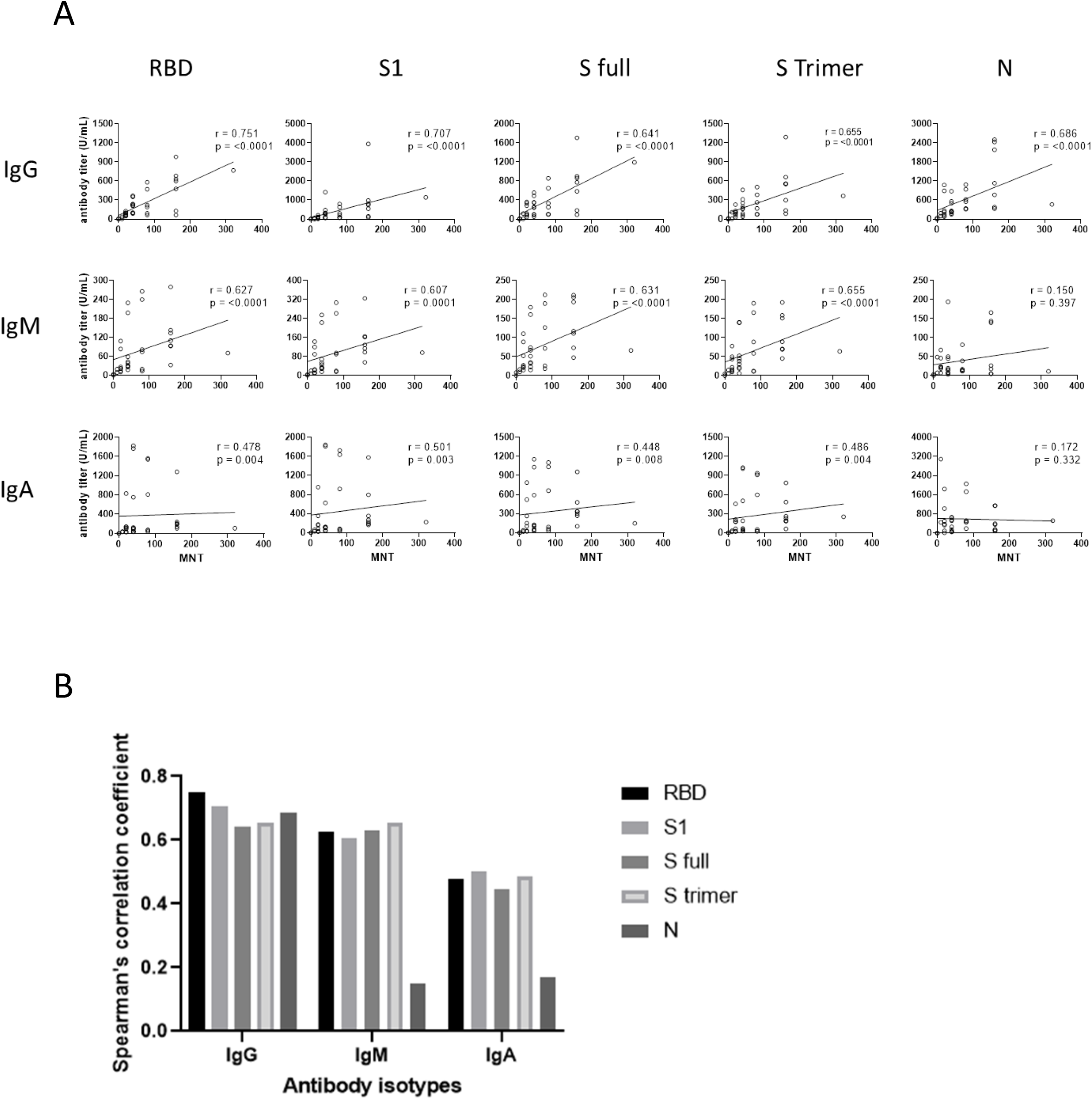
Correlation between antigen-specific antibody isotypes and neutralizing activity. (A) Correlation analysis between antibody titre for each SARS-CoV-2 antigen and microneutralization titre was calculated using Spearman’s correlation coefficient. Serological and neutralization assays were performed using 34 serums from 19 patients. (B) Graph represents the Spearman’s coefficient (r) in (A).

### Relationship between RBD-IgG production and disease severity

We categorized the COVID-19 patients into two severity groups (severe and non-severe) based on established clinical classifications. The average levels of RBD-IgG in the serum samples, which was the best indicator of neutralizing activity according to Fig. 3, are shown in Fig. 4. Serum RBD-IgG levels in severe COVID-19 patients were significantly higher than non-severe cases as early as 8–14 days after symptom onset, while severe COVID-19 patients produced high levels of RBD-IgG over time.

**Figure 4.**
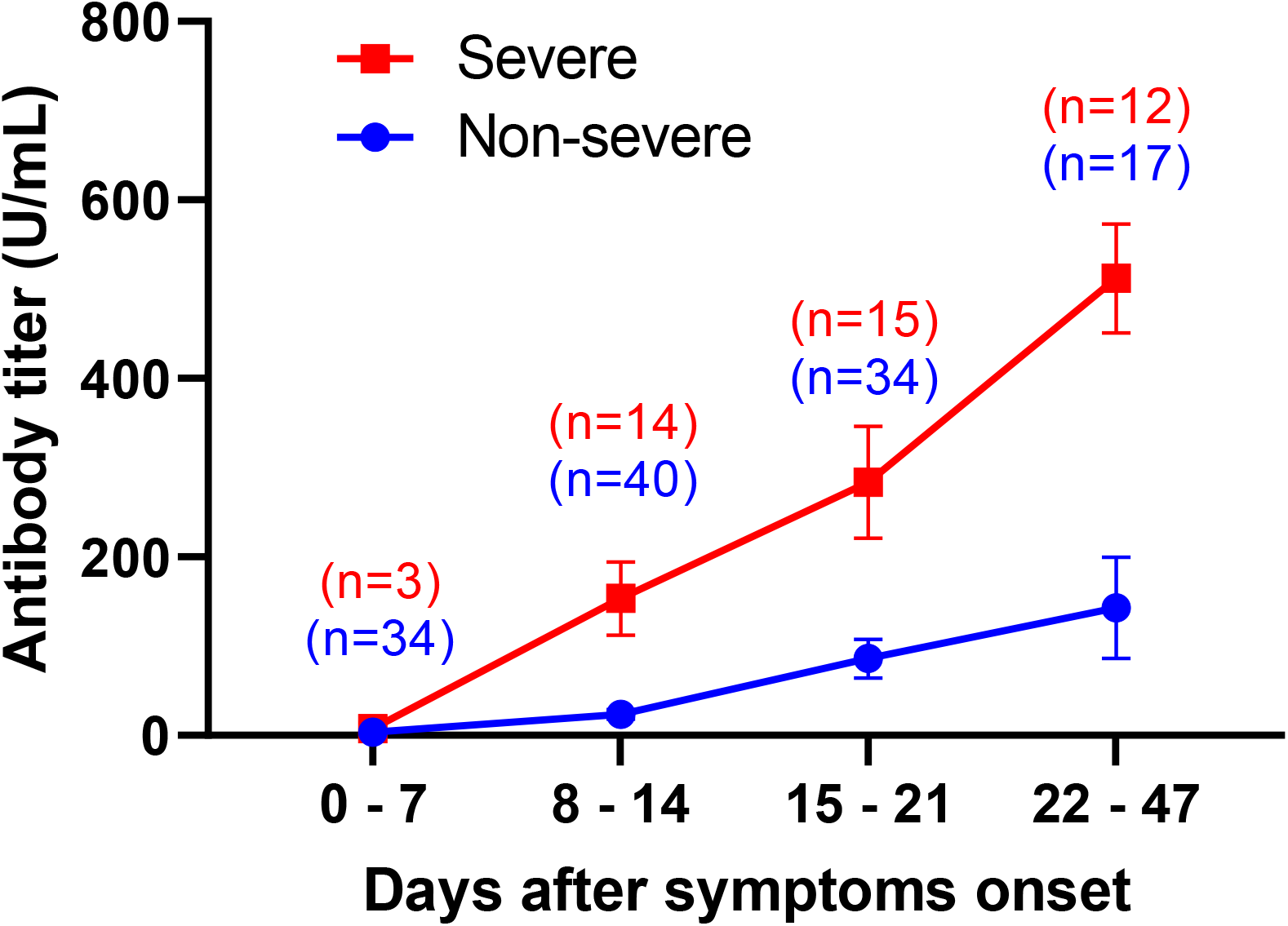
Relationship between IgG antibody titre against RBD and disease severity. IgG antibody titre against RBD (RBD-IgG) were classified into two severity groups: severe and non-severe. Red is severe (6 patients) and blue is non-severe (35 patients). Statistical analysis was done by two-tailed Mann-Whitney test (** P < 0.01, *** P <0.001).

### Kinetics of serum RBD-IgG levels and neutralizing activity in severe COVID-19 patients

We evaluated the relationship between RBD-IgG responses and virus neutralizing activity in serum samples obtained from four severe COVID-19 patients at several time points after symptom onset. They all recovered from COVID-19 and duration of the COVID-19-associated symptoms was 21, 23, 22 and 37 for patients 5, 14, 31 and 34, respectively. As shown in Fig. 5, serum RBD-IgG in all four patients increased over time, in concordance with virus neutralizing activity.

**Figure 5.**
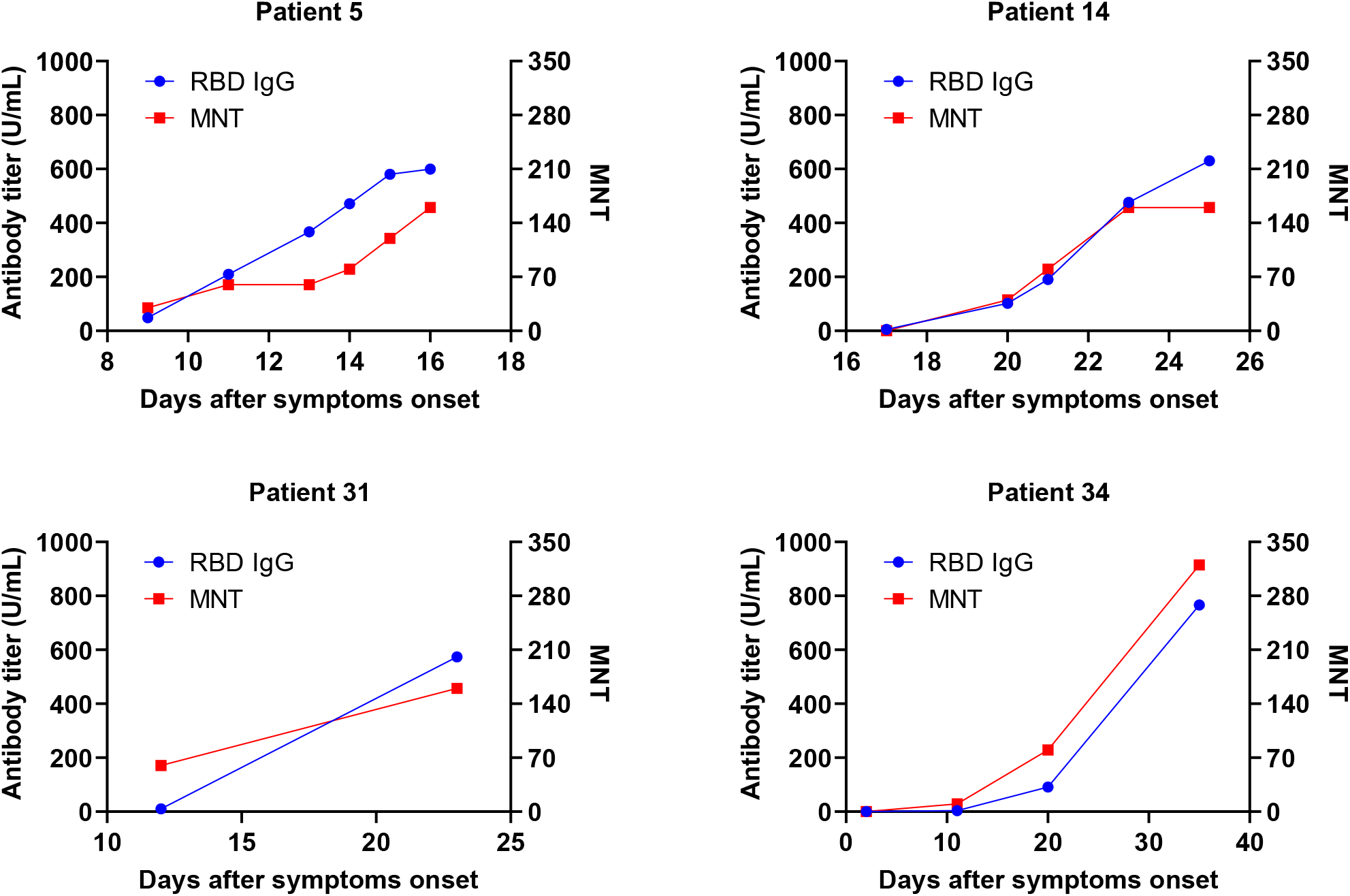
Kinetics of RBD-IgG antibody titers and neutralizing activity in patients with severe COVID-19. Relationship between RBD-IgG titers and microneutralization titers (MNT) in serum samples from four patients with severe COVID-19 over several time periods after symptom onset.

## Discussion

In this study, we comprehensively analyzed antigen-specific anti-SARS-CoV-2 antibody isotypes, including IgG, IgM and IgA in COVID-19 patients and identified differential antibody responses and their relationship with neutralizing activity among the isotypes. The insights into the reactivity of each antibody isotype against various antigens generated here should further improve our understanding of the humoral immune responses against SARS-CoV-2.

We investigated the kinetics of RBD, S1, S full, S trimer and N protein specific IgG, IgM and IgA responses in 169 serum samples obtained from 41 COVID-19 patients. Our study showed that antigen-specific antibody isotypes in the sera from several patients increased within 0–7 days after the onset of symptoms (Fig. 1 and Table 2). The average of all antigen-specific IgA levels reached their peaks in 15–21 days after the onset of symptoms, while all antigen-specific IgG and IgM levels significantly increased in 8–14 days after the onset and continued to increase thereafter. These increases in IgG and IgM are consistent with some recent reports (20-24). Although there are few available reports on the responses of IgA in COVID-19 patients, one report showed that RBD-specific IgA reached the peak during 16–20 days after the onset of symptoms, which is consistent with our findings (22). These results indicate that, although there are some differences among the isotypes, all antigen-specific antibody isotypes can be detected in the serum of most patients about 2 weeks after the onset of symptoms.

However, our further analysis revealed that there are considerable differences among the antibodies in terms of the specificity and sensitivity in predicting disease. When the specificity of each antigen-specific antibody isotypes was investigated using a cut-off value determined by the ROC analysis, specificity of the S protein assay for all isotypes showed high specificities ranging between 94.0%–99.0%, while the N protein assay for all isotypes showed lower specificities of 79.0%, 88.0% and 82.0% for N-IgG, N-IgM and N-IgA, respectively. The lower specificity of N protein may be due to potential cross-reactivity with antibodies to commonly circulating CoVs in the negative control samples. Although we have not addressed the potential cross-reactivity of each antigen with serum from humans infected with other CoVs, several studies have shown that N protein-based serological assays were more often associated with cross-reactivity than S protein-based assays (25). Chia et al. reported that S1 or RBD showed better specificity than N-based serological assay and that there was significant cross-reactivity when N protein was used as the antigen for their assay (26). Furthermore, Algaissi et al. reported that the specificity of N-based IgG and IgM assays showed lower sensitivity than S1-based assays (23). Our findings support the use of S protein as the antigen for the detection of SARS-CoV-2 specific antibodies.

We also determined which antigen-specific antibody isotype assays best represented the virus neutralizing activity (Fig. 3A and B). Although most of the antibody isotypes showed good correlation with neutralizing activity, RBD-IgG assay showed the best correlation. These results are in line with the findings of other reports, which showed correlation of RBD-IgG levels with neutralizing activity (27, 28). Neutralization assays are difficult to perform as routine tests as they require viral cultures and must be performed in laboratories with higher biosafety levels. Measurement of RBD-IgG levels in the serum can be a reliable and convenient tool for assessing the immunological response of COVID-19 patients. We also suggest that measuring RBD-IgG levels in the serum may be a useful tool to identify donors for convalescent serum/plasma therapy and quantify the immunogenicity of vaccines which are currently being developed (10, 29, 30).

Since RBD-IgG levels in the serum correlated with virus neutralizing activity, we investigated the relationship between severity of COVID-19 patients and serum RBD-IgG levels. As shown in Fig. 4, RBD-IgG levels in the serum samples collected from severe COVID-19 patients within 8–14 days after symptom onset were significantly higher than those of non-severe patients; however, the severe COVID-19 patients showed much higher levels of RBD-IgG thereafter. Consistent with previous reports, our results indicate that measurement of RBD-IgG levels correlate with disease severity of COVID-19 (22, 31). Furthermore, the serum RBD-IgG levels in severe COVID-19 patients who recovered increased along with neutralizing activity. These results suggest that measurement of RBD-IgG levels in convalescent patients can be used to identify appropriate donors with high neutralizing activity for convalescent serum/plasma therapy (7, 10).

There are several limitations in this study. Importantly, there were only four severe COVID-19 patients from whom residual serum samples were available to investigate the association between serum RBD-IgG levels and neutralizing activity. Since these four severe COVID-19 patients all recovered from COVID-19, we could not investigate differences in antibody responses and neutralizing activity between patients who recovered from COVID-19 and those who did not.

In summary, our results indicate that the anti-SARS-CoV-2 antibody response in COVID-19 patients varies among the antigen-specific antibody isotypes. Diagnostic performance of ELISAs for detection of anti-SARS-CoV-2 antibodies also varies among antigen-specific antibody isotypes. Among them, serum RBD-IgG levels best correlate with virus neutralizing activity and disease severity, thus may be the optimal assay to track COVID-19 seroconversion responses and use as the basis for COVID-19 serological tests.

## Acknowledgments

We would like to thank Dr. Hideyuki Saya (Keio University School of Medicine) for the critical discussions and suggestions.

